# The Synthetic Fidelity–Stability Framework (SFSF): A Systematic Multi-Dimensional Benchmark of Synthetic Clinical Laboratory Data Generators

**DOI:** 10.64898/2026.08.11.741471

**Authors:** Sachin S Desh, Prakash MG Achary, Saurav Nayak

**Affiliations:** ICMR National Institute of Child Health Research, Ansari Nagar West, New Delhi, India; Department of Biochemistry, All India Institute of Medical Sciences, New Delhi,India

## Abstract

**Background:** Synthetic data generation is increasingly proposed as a strategy to support privacy-preserving data sharing, augmentation of small or restricted biomedical datasets, and benchmarking of artificial intelligence tools in laboratory medicine. However, model selection remains difficult because synthetic data generators differ in fidelity, privacy risk, stability, and generalisability. Existing evaluations have rarely examined performance jointly across conditioning signal strength, synthetic output scale, and train–test generalisation.

**Methods:** We developed the Synthetic Fidelity–Stability Framework (SFSF), a systematic benchmark of 17 synthetic tabular data generation models using NHANES as a complex biomedical reference dataset. Models included statistical, copula-based, resampling, variational autoencoder, generative adversarial network, and diffusion-based approaches. Synthetic datasets were generated across 11 seed sizes, from 0 to 500 real conditioning observations, and six output scales, from 50 to 5,000 rows, yielding 1,122 synthetic datasets per run. Each dataset was evaluated against the full original dataset, the training subset, and a held-out test subset across five tiers: univariate distributional fidelity, moment agreement, tail behaviour, multivariate dependency structure, and privacy/memorisation risk. Composite rankings and seed-versus-output stability profiles were derived.

**Results:** Univariate fidelity was broadly recovered across model classes and was the least discriminating tier. Resampling-based methods ranked highest overall but showed the greatest privacy risk, reflecting proximity to real observations rather than true generative novelty. VAE-family models reproduced moment statistics relatively well but consistently failed on tail and shape fidelity. GAN-family models showed substantial moment-level instability, while VineCopula demonstrated severe multivariate dependency failure. Diffusion-based models, particularly ForestDiffusion, provided the most favourable privacy–utility balance, combining competitive fidelity with the lowest privacy risk and the smallest train–test gap.

**Conclusions:** No single synthetic data generator dominated across fidelity, stability, and privacy dimensions. The SFSF framework provides a practical, multi-criterion approach for selecting synthetic tabular data generators according to intended clinical laboratory use, balancing statistical realism, dependency preservation, privacy risk, and robustness to seed and output scale.

## Introduction

Access to large, representative clinical laboratory datasets is increasingly essential for developing, validating, and benchmarking artificial intelligence and machine learning tools in laboratory medicine (1,2). However, patient privacy regulations and institutional data governance requirements often restrict inter-laboratory data exchange, constrain dataset size, and limit collaborative evaluation of AI-driven laboratory algorithms (2–4). Niche medical datasets are often small, institution-specific, sparsely annotated, and constrained by privacy or governance restrictions, as in the case of specific diseases, outbreaks or other sparse conditions, making conventional big-data AI/ML development difficult or impossible (5,6). This creates a critical lacuna in laboratory medicine, where clinically relevant modelling problems frequently lack the scale, diversity, and shareability of data needed for robust algorithm development and external validation. Thus, there exists a persistent structural tension between the scale of data required for robust model development and validation, and the volume and diversity of data that can realistically be accessed, shared, or reused in practice (7).

Synthetic data generation (SDG), defined as the computational creation of artificial datasets that approximate the statistical properties of real clinical data without directly containing individual patient records, has emerged as a promising strategy to address this tension (7,8). By separating analytical utility from direct access to identifiable patient-level data, SDG may support privacy-preserving data sharing, augmentation of small or imbalanced datasets, inter-laboratory collaboration, and the development of training datasets for diagnostic AI systems (2,9,10). In clinical laboratory medicine, proposed and demonstrated applications include reference interval derivation, method validation, patient-based real-time quality control (PBRTQC), proficiency testing and external quality assessment (EQA) design, and educational case generation (2,11,12). A diverse set of generative approaches has been applied to tabular health data, including classical statistical models, copula-based methods, variational autoencoders (VAEs), generative adversarial networks (GANs), and, more recently, diffusion-based models (2,12,13). These model classes differ substantially in their assumptions about data structure and generation mechanisms, with important implications for statistical fidelity, privacy risk, and robustness under different deployment conditions (12,14).

Despite increasing interest in SDG, the evidence base for selecting appropriate models in biomedical tabular data remains incomplete. Prior benchmark studies have shown that no universal evaluation strategy currently exists, that generalisability across datasets remains limited and that no single model consistently outperforms others across utility, fidelity, and privacy criteria (15). Consequently, model selection must remain context-specific and guided by the intended clinical or analytical use case. Yan et al. further demonstrated that the privacy–utility trade-off varies considerably across applications, while inconsistent terminology and heterogeneous evaluation criteria continue to limit standardised quality assessment across the literature (16,17). Importantly, no published study has systematically examined how SDG model performance varies jointly with conditioning signal strength, the amount of real data used to seed generation, and synthetic output scale. These factors are highly relevant to laboratory practice, where available real data may range from small reference cohorts to thousands of historical observations, and where synthetic output requirements may span compact validation datasets to large-scale simulation populations. In addition, existing evaluation frameworks have not consistently separated in-sample reconstruction from out-of-sample generalisation in broad model comparisons. This distinction is clinically important: a generator that reproduces training-set structure rather than learning the underlying data-generating distribution offers limited generalisable utility and may also carry increased privacy risk (16,17).

The present study addresses these gaps through the **Synthetic Fidelity–Stability Framework (SFSF)**, a systematic, multi-factorial benchmark of 17 SDG models applied to the National Health and Nutrition Examination Survey (NHANES), a complex, high-dimensional biomedical tabular reference dataset (18). It evaluates model performance across five quality tiers: univariate distributional fidelity, moment agreement, tail behaviour, multivariate dependency structure, and privacy risk.

## Materials and Methods

### 2.1 Study Design

This study, as elucidated in Figure 1, was designed as a systematic, multi-factorial benchmark experiment evaluating multiple (n=17) synthetic data generation models across controlled variations in conditioning signal strength and generation scale. The experimental framework was designed to simultaneously assess fidelity, statistical validity, structural integrity, and privacy risk, while explicitly distinguishing in-sample reconstruction from out-of-sample generalisation. No patient data were collected for this study. NHANES data are publicly available and exempt from ethical review requirements; no institutional ethics approval was required.

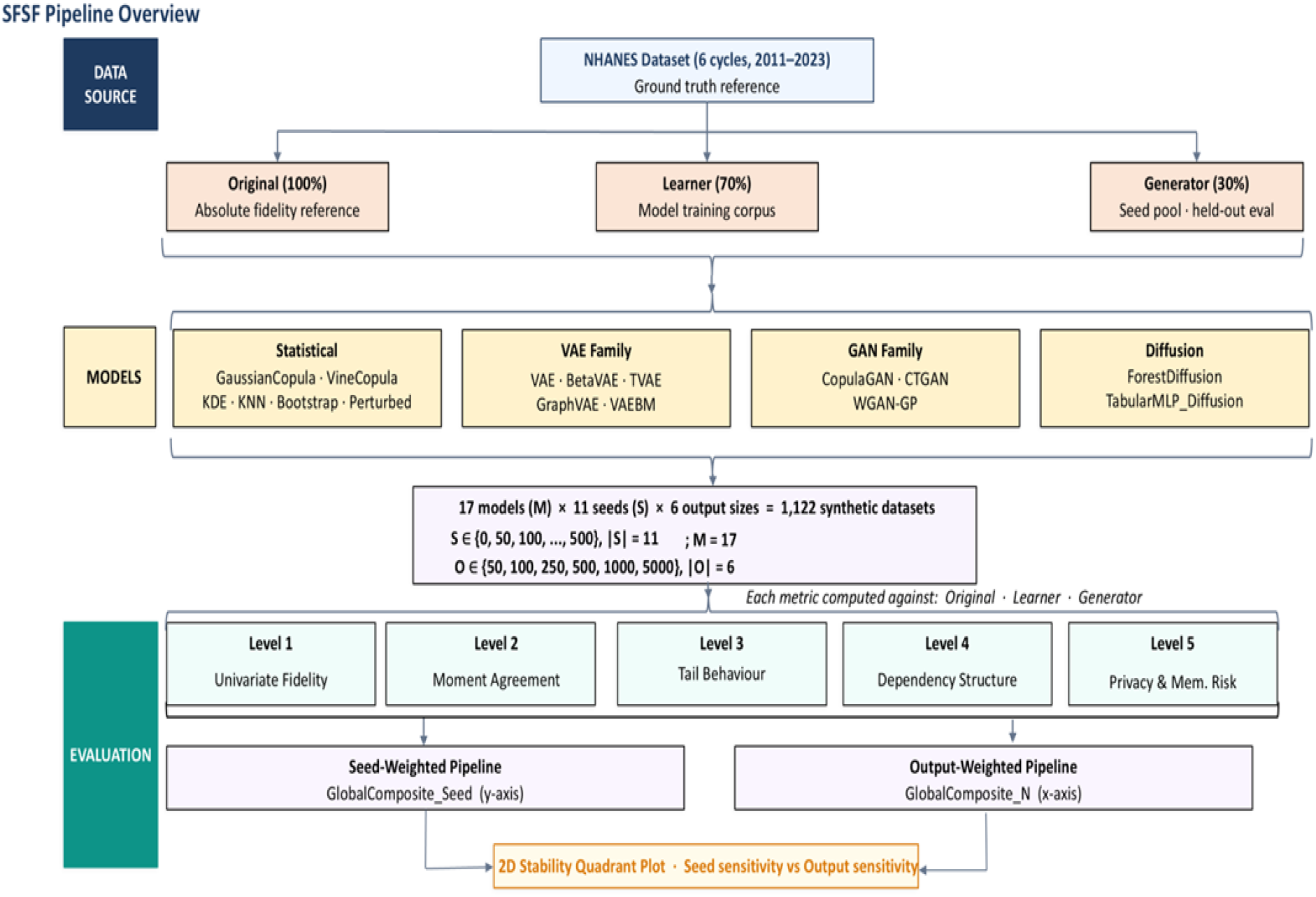

### 2.2 Reference Dataset

NHANES is a continuous cross-sectional programme conducted by the Centers for Disease Control and Prevention’s National Center for Health Statistics, and was used as the ground truth reference dataset. NHANES combines standardised physical examination, clinical laboratory testing, and questionnaire data in a nationally representative sample of the United States civilian population, producing a complex, high-dimensional dataset encompassing continuous laboratory analytes, categorical demographic variables, and mixed-type clinical measurements [16,17]. These structural characteristics — including inter-analyte correlations, non-normal marginal distributions, and mixed variable types — are representative of those encountered in clinical laboratory information systems, making NHANES an established and appropriate benchmark for tabular SDG evaluation [8,9]. The NHANES cycles used The NHANES cycles utilised were 2011–2012, 2013–2014, 2015–2016, 2017– 2018, the 2017–March 2020 pre-pandemic combined release, and 2021–2023 [15,16]. The 2017–March 2020 dataset represents the expanded combined release incorporating the 2017–2018 cycle data alongside additional pre-pandemic collection; both were retained as distinct cycle-level datasets to preserve temporal granularity across the full analytical period.

### 2.3 Dataset Partitioning

The full dataset was partitioned into three disjoint subsets with distinct analytical roles. The Original dataset (100%) served as the absolute ground truth reference for fidelity assessment. The Learner subset (70%) constituted the exclusive training corpus available to all SDG models. The Generator subset (30%) was withheld entirely from model training and used solely for seed construction and out-of-sample evaluation. This tripartite design enables simultaneous assessment of in-sample reconstruction and out-of-sample generalisation, with divergence between Learner-referenced and Generator-referenced evaluation scores serving as a model-level index of overfitting (17,19).

### 2.4 Synthetic Data Generation Models

Seventeen SDG models were evaluated, spanning four methodological classes. Classical and statistical models included GaussianCopula (GC), VineCopula (VC), IndependentMarginal (IM), Kernel Density Estimation (KDE), k-Nearest Neighbours resampling (KNN), JitteredBootstrap (JB), and PerturbedReal (PR). Variational autoencoder family models included VAE, β-VAE (BetaVAE), Tabular VAE (TVAE), GraphVAE, and VAEBM.

Generative adversarial network models included CopulaGAN, Conditional Tabular GAN (CTGAN), and Wasserstein GAN with gradient penalty (WGAN-GP). Diffusion-based models included ForestDiffusion (FD) and TabularMLP_Diffusion (TMD) (13). All models were implemented using Python 3.1.1 and executed under consistent computational conditions to ensure that performance differences reflect model characteristics rather than experimental variability.

### 2.5 Experimental Design

Synthetic data generation was conditioned on 11 seed datasets, each drawn from the held-out Generator subset. Seed sizes ranged from 0 to 500 real observations in increments of 50. The zero-seed condition represents a cold-start generative regime in which models rely exclusively on learned priors without real-data conditioning; increasing seed sizes progressively inject real observations, enabling assessment of model sensitivity to conditioning strength. For each seed configuration, synthetic datasets were generated at six predefined output scales: 50, 100, 250, 500, 1,000, and 5,000 rows. This range was selected to probe model behaviour across low-sample and high-sample generation regimes relevant to laboratory applications. The complete factorial design of 17 models × 11 seed configurations × 6 output scales yielded 1,122 synthetic datasets per experimental run. Datasets were stored with a standardised naming convention encoding model type, seed size, and output size to ensure traceability and reproducibility.

### 2.6 Evaluation Framework

Model performance was assessed using a hierarchical five-tier metric framework. Level 1 (Univariate Distributional Fidelity) evaluated per-variable marginal distribution agreement using the Kolmogorov–Smirnov statistic, Jensen–Shannon divergence, Wasserstein distance, and coverage of the real value support (20–23). Level 2 (Moment and Summary Statistic Agreement) quantified deviations in central tendency and dispersion through standardised mean difference, variance ratio, skewness difference, and median absolute error (12). Level 3 (Tail Behaviour and Distribution Shape) assessed kurtosis difference, 99th percentile error, outlier rate parity, and upper tail dependence — metrics of particular relevance to laboratory reference interval modelling and risk stratification (12). Level 4 (Multivariate Dependency Structure) evaluated preservation of inter-variable relationships through pairwise Pearson correlation difference, Kendall’s τ difference, correlation matrix distance, and contingency similarity for categorical variable pairs (24,25). Level 5 (Privacy and Memorisation Risk) quantified proximity of synthetic records to real observations using Distance to Closest Record (DCR), Nearest-Neighbour Distance Ratio (NNDR), exact match share, and membership inference attack (MIA) success rate (16,19). All metrics were computed against each of the three reference datasets — Original, Learner, and Generator — yielding 15 composite scores per model prior to aggregation.

### 2.7 Composite Metric Construction and Stability Analysis

Sub-metrics within each evaluation tier were aggregated into per-level composite scores computed separately for each reference dataset. Two parallel aggregation pipelines were constructed: a seed-weighted pipeline, aggregating across output sizes to produce a seed-conditioned global composite (GlobalComposite-Seed); and an output-weighted pipeline, aggregating across seed configurations to produce an output-conditioned global composite (GlobalComposite-N). Final model rankings were derived using scalar ranking by global composite score, Pareto dominance ranking across all five levels (Scalar Rank), and a combined final rank integrating both approaches (Meta Rank). To characterise model robustness, GlobalComposite-Seed and GlobalComposite-N scores were jointly plotted, partitioning models into four stability quadrants at the cross-model mean: robust on both axes; seed-sensitive and output-stable; output-sensitive and seed-stable; and sensitive on both axes.

## Results

Seventeen synthetic data generation models were evaluated across 1,122 unique experimental conditions defined by 11 seed configurations (0–500 real rows) and 6 synthetic output sizes (50–5,000 rows), with NHANES serving as the ground truth reference. Each synthetic dataset was assessed against three reference distributions—Original (100%), Learner (70%), and Generator (30% held-out)—across five evaluation tiers spanning univariate fidelity (Level 1), moment agreement (Level 2), tail and shape fidelity (Level 3), multivariate dependency structure (Level , and privacy risk (Level 5). Global composites were derived through two parallel pipelines—seed-weighted (S-weighted) and output-size-weighted (N-weighted)—and aggregated into a meta-ranking. The S-weighted pipeline aggregates performance across output sizes for each seed level, while the N-weighted pipeline aggregates across seed configurations for each output size.

### Level 1: Univariate Distribution Fidelity

All models achieve low composite error (range 0.002–0.045), indicating that marginal distributions are broadly recovered regardless of architecture. GaussianCopula and KNN return the lowest L1 scores (0.003), while VineCopula is the sole clear outlier (0.045) in the N-weighted pipeline. Resampling methods, diffusion models, and the VAE family cluster within 0.003–0.010, suggesting that univariate coverage is not architecturally limiting for any class of model tested.

### Level 2: Moment & Summary Statistics Agreement

The GAN family produces the worst moment agreement by a substantial margin: CTGAN scores 0.587, CopulaGAN 0.500, and WGAN_GP 0.361—collectively more than twice the mean of the resampling family (0.182) and more than four times the VAE-family mean (0.114). The VAE family achieves the lowest mean L2 score (0.114), with BetaVAE (0.073) performing comparably to PerturbedReal (0.137). The disentanglement constraint in BetaVAE appears to impose a regularisation effect that stabilises moment reproduction. Diffusion models score moderately (ForestDiffusion 0.259, TabularMLP_Diffusion 0.306), suggesting that iterative refinement partially, but not fully, corrects moment-level errors introduced during the diffusion process.

### Level 3: Tail & Shape Fidelity

The VAE family fails uniformly on tail and shape fidelity (scores 0.313–0.347), indicating that while VAEs reproduce moment statistics well, they systematically smooth extreme values and distort distributional shape—a consequence of latent Gaussian assumptions and KL-regularised encoding. Notably, TabularMLP_Diffusion (L3 = 0.352) performs as poorly as the worst VAE-family models despite its superior L4 score, suggesting that MLP-based diffusion models share the VAE pathology of extremal smoothing. KDE achieves the best L3 score overall (0.070 seed-weighted), followed by JitteredBootstrap (0.101) and KNN (0.134). The strong L3 performance of resampling methods is structurally expected: copying or lightly perturbing real rows preserves tail structure by construction. CopulaGAN (0.088) is the best-performing generative model on L3, surpassing all GAN and diffusion peers.

### Level 4: Multivariate Dependency Structure

Resampling methods and GaussianCopula achieve the best L4 scores (0.055–0.071), reflecting their reliance on observed joint structure rather than modelled decompositions. Among fully generative models, TabularMLP_Diffusion (0.055) and WGAN_GP (0.056) match resampling-level dependency preservation—a remarkable result that underscores the capacity of these architectures to recover inter-column correlation structure.

### Level 5: Privacy & Memorisation Risk

Resampling methods—which dominate the composite ranking—exhibit the highest privacy risk: JitteredBootstrap (0.245), PerturbedReal (0.264), and KNN (0.255) all return elevated L5 scores driven by small DCR, elevated NNDR, and non-trivial exact-match rates. The two diffusion models represent the most favourable privacy-utility balance: ForestDiffusion (L5 = 0.098) and TabularMLP_Diffusion (L5 = 0.099) achieve the lowest privacy risk scores of any model class while maintaining competitive L1–L4 performance. GaussianCopula (0.185) and VineCopula (0.127) also perform well on L5, though the latter’s L4 failure significantly limits practical utility. Among VAE models, VAEBM (0.171) has marginally elevated risk relative to its peers, while BetaVAE (0.164) and TVAE (0.161) are comparable.

### Global Model Ranking

Table 1 presents per-level scores (N-weighted) alongside global composite and meta-rank for all 17 models, sorted by meta-rank.

**Table 1:** Global model ranking. Per-level N-weighted composite scores and meta-rank (lower = better)

| Model | Level 1 | Level 2 | Level 3 | Level 4 | Level 5 | Global | Scalar Rank | Meta Rank |
| --- | --- | --- | --- | --- | --- | --- | --- | --- |
| PerturbedReal | 0.003 | 0.137 | 0.117 | 0.067 | 0.264 | **0.061** | 2 | 1.5 |
| JitteredBootstrap | 0.003 | 0.140 | 0.101 | 0.067 | 0.245 | **0.058** | 1 | 2.0 |
| KNN | 0.002 | 0.204 | 0.134 | 0.070 | 0.255 | **0.065** | 3 | 2.0 |
| GaussianCopula | 0.003 | 0.333 | 0.156 | 0.071 | 0.185 | **0.072** | 5 | 2.75 |
| ForestDiffusion | 0.004 | 0.269 | 0.153 | 0.090 | 0.098 | **0.069** | 4 | 3.0 |
| TabularMLP_Diffusion | 0.005 | 0.306 | 0.352 | 0.055 | 0.099 | **0.078** | 6 | 3.75 |
| KDE | 0.006 | 0.266 | 0.200 | 0.140 | 0.291 | **0.105** | 11 | 3.75 |
| CopulaGAN | 0.006 | 0.500 | 0.088 | 0.132 | 0.202 | **0.094** | 7 | 5.0 |
| BetaVAE | 0.010 | 0.157 | 0.313 | 0.159 | 0.164 | **0.104** | 10 | 5.25 |
| CTGAN | 0.008 | 0.587 | 0.098 | 0.137 | 0.203 | **0.106** | 13 | 6.5 |
| GraphVAE | 0.009 | 0.153 | 0.313 | 0.155 | 0.159 | **0.102** | 8 | 6.5 |
| WGAN_GP | 0.029 | 0.361 | 0.223 | 0.056 | 0.130 | **0.111** | 16 | 6.5 |
| VAE | 0.010 | 0.148 | 0.315 | 0.152 | 0.164 | **0.102** | 8 | 6.75 |
| TVAE | 0.009 | 0.163 | 0.316 | 0.164 | 0.161 | **0.105** | 11 | 6.75 |
| VAEBM | 0.010 | 0.119 | 0.323 | 0.221 | 0.171 | **0.108** | 15 | 7.5 |
| VineCopula | 0.045 | 0.289 | 0.145 | 0.641 | 0.131 | **0.173** | 17 | 7.75 |
| IndependentMarginal | 0.007 | 0.270 | 0.173 | 0.140 | 0.291 | **0.107** | 14 | 8.25 |

Three resampling-based methods—PerturbedReal (MetaRank 1.5), JitteredBootstrap (2.0), and KNN (2.0)— occupy the top positions, followed by GaussianCopula (2.75) and ForestDiffusion (3.0). The supremacy of resampling methods reflects their structural advantage: by closely mirroring or lightly perturbing real data, they achieve near-zero composite error at the cost of limited generative novelty. This is a necessary caveat for practical interpretation—superiority in composite rank does not imply suitability for privacy-sensitive deployment, as discussed in Section 5.

The five VAE-family models (VAE, GraphVAE, TVAE, VAEBM, BetaVAE) rank in the 5th–8th tier of the meta-ranking, forming a compact cluster with global composites in the range 0.102–0.116. GAN-family models are split: CopulaGAN (5.0) performs comparably to the upper VAE tier, whereas CTGAN (6.5) and WGAN_GP (6.5) show higher composite error driven by level-specific failures. VineCopula (7.75) and IndependentMarginal (8.25) consistently rank last. Table 2 summarises mean per-level composite scores by model family, enabling direct comparison of failure modes across architectural paradigms.

**Table 2:** Mean per-level N-weighted composite scores by model family.

| Model Family | n | Level 1 | Level 2 | Level 3 | Level 4 | Level 5 |
| --- | --- | --- | --- | --- | --- | --- |
| Resampling | 3 | 0.003 | 0.182 | 0.117 | 0.068 | 0.258 |
| Statistical-Copula | 2 | 0.003 | 0.269 | 0.162 | 0.356 | 0.152 |
| Diffusion | 2 | 0.005 | 0.288 | 0.253 | 0.073 | 0.099 |
| VAE family | 5 | 0.010 | 0.114 | 0.319 | 0.240 | 0.159 |
| GAN family | 3 | 0.014 | 0.483 | 0.136 | 0.109 | 0.179 |
| Independent Marginal | 1 | 0.007 | 0.270 | 0.173 | 0.140 | 0.291 |
| KDE | 1 | 0.006 | 0.266 | 0.200 | 0.140 | 0.291 |

### Overfitting Diagnostics: Train-Test Gap

Among all models, ForestDiffusion exhibits the smallest mean train-test gap across Levels 1–4 (−0.003), indicating near-perfect generalisation. In contrast, TabularMLP_Diffusion (−0.031), VAE (−0.024), GaussianCopula (−0.022), and WGAN_GP (−0.023) show the largest gaps, signalling meaningful overfitting to the training partition. The resampling methods show small to moderate gaps (−0.005 to −0.010) despite their elevated L5 scores, consistent with the interpretation that their privacy risk arises from structural copying rather than distributional overfitting.

### Seed & Output – Scaled Sensitivity

Positioning models on a two-dimensional sensitivity space—GlobalComposite-Seed (y-axis) against GlobalComposite-N (x-axis), with quadrant boundaries at cross-model means—reveals four behavioural regimes summarised in Figure 1.

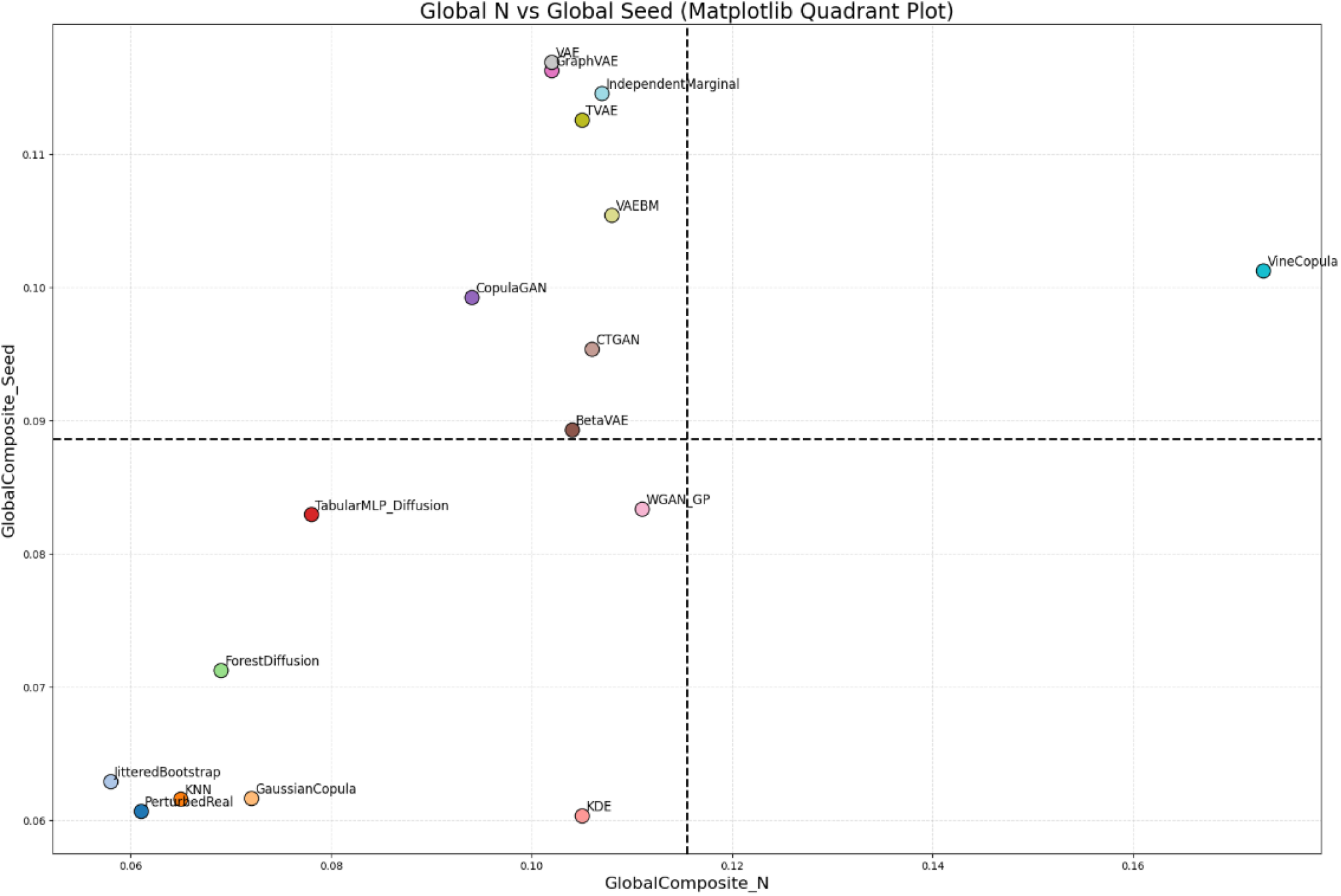

## Discussion

The benchmarking results of the Synthetic Fidelity–Stability Framework reveal critical performance disparities across the 17 generative architectures, particularly between legacy statistical methods, Variational Autoencoders, Generative Adversarial Networks, and emerging Diffusion models. Consistent with recent comparative studies, while GAN-based models like CTGAN and MedGAN have historically set the standard for synthetic Electronic Health Record generation, they frequently suffer from training instability and mode collapse, which limits their ability to capture the full diversity of clinical laboratory distributions (26,27). In contrast, diffusion-based approaches, such as TabDDPM, demonstrate a superior capacity to generate realistic mixed-type tabular data by leveraging iterative denoising processes that provide more stable training dynamics and higher fidelity across complex clinical features (28,29). The SFSF Global Model Ranking highlights that while no single model excels in every dimension, the shift toward diffusion and hybrid graph-guided architectures often yields a more reliable “integrity frontier” for clinical applications (30,31).

A pivotal finding in the evaluation of univariate fidelity and tail behavior is the varying ability of models to represent “outlier” or pathological lab values. In clinical laboratory medicine, abnormal results in the distribution tails are often more diagnostically significant than central tendencies (32). Our results indicate that many traditional models struggle with these extremes, potentially leading to “distributional bias” where rare but critical clinical states are underrepresented. Recent evidence suggests that Forest Diffusion and specific Denoising Diffusion Probabilistic Models are significantly more adept at capturing these tail characteristics compared to standard Gaussian Copulas or CTGAN (32,33). This suggests that for high-stakes laboratory validation and method comparison, researchers must prioritize models that exhibit high “Tail Fidelity” to ensure that synthetic cohorts remain clinically valid for stress-testing diagnostic algorithms (2,34).

The analysis of multivariate dependency further underscores the complexity of capturing the inherent logical relationships between different laboratory measurands, such as the correlation between creatinine levels and estimated glomerular filtration rate (eGFR). Models that neglect the local neighborhood structure or the manifold geometry of the data often fail to preserve these pairwise and high-order correlations (31). The performance of tree-based ensembles and graph-guided latent diffusion models in our benchmark confirms that integrating structural priors—such as patient similarity graphs—can reduce marginal distribution errors and improve the coherence of joint feature structures (30,31). This multivariate integrity is essential for downstream machine learning utility, as synthetic data must not only look realistic in isolation but also maintain the predictive relationships required to train robust clinical decision support systems (35,36).

Privacy and memorization risk remain the most significant barriers to the widespread adoption of synthetic clinical data. Our framework identifies a clear utility–privacy trade-off: models achieving the highest fidelity scores often exhibit the highest risk of “identity disclosure” or “membership inference,” suggesting they may be memorizing specific training exemplars rather than learning generalizable patterns (29,37). While recent frameworks like EHR-Safe and SynQP have introduced rigorous privacy audits—including Distance to Closest Record and Nearest Neighbor Adversarial Accuracy—many generative models still lack native differential privacy guarantees (36,38). The benchmarking of the “Train-Test Gap” in SFSF serves as a vital overfitting diagnostic, warning against the deployment of models that appear high-performing but effectively “leak” sensitive patient information through near-verbatim replication of the source data (2,17).

The stability dimension of the SFSF framework—specifically the sensitivity to random seeds and output scaling— addresses a frequently overlooked aspect of synthetic data generation: reproducibility. Clinical laboratories require “Calibration Stability” to ensure that the synthetic data used for method validation or quality control does not vary wildly between training runs (2,30). Our findings suggest that diffusion models and certain tree-based methods offer more consistent performance across different initialization seeds compared to GANs, which are notoriously sensitive to hyperparameter configurations (27,28). This stability is not merely a technical preference but a requirement for “trustworthy AI” in healthcare, where the reliability of a synthetic dataset must be guaranteed before it can be used for regulatory submissions or external quality assessments (30,39).

The implementation of synthetic data in the clinical laboratory offers a transformative path for democratizing data access, augmenting rare disease cohorts, and validating AI workflows without compromising patient confidentiality (2). However, the SFSF results suggest that implementation must be guided by a multi-dimensional “RSCE” perspective rather than simple fidelity metrics (30). As regulatory frameworks evolve, the use of hybrid real-synthetic datasets and the adoption of standardized benchmarking ontologies will be essential for moving synthetic data from experimental “sandboxes” into routine laboratory practice (2,40). Future research should focus on refining multimodal generation to integrate longitudinal lab results with clinical notes and medical imaging, creating a more holistic synthetic patient record for the next generation of precision medicine (35,41).

## Data Availability Statement

The datasets used and/or analysed during the current study available from the corresponding author on reasonable request.

